# Improving the visualisation, interpretation and analysis of two-sample summary data Mendelian randomization via the radial plot and radial regression

**DOI:** 10.1101/200378

**Authors:** Jack Bowden, Wesley Spiller, Fabiola Del Greco-M F, Nuala Sheehan, John Thompson, Cosetta Minelli, George Davey Smith

## Abstract

**Background:** Summary data furnishing a two-sample Mendelian randomization study are often visualized with the aid of a scatter plot, in which single nucleotide polymorphism (SNP)-outcome associations are plotted against the SNP-exposure associations to provide an immediate picture of the causal effect estimate for each individual variant. It is also convenient to overlay the standard inverse variance weighted (IVW) estimate of causal effect as a fitted slope, to see whether an individual SNP provides evidence that supports, or conflicts with, the overall consensus. Unfortunately, the traditional scatter plot is not the most appropriate means to achieve this aim whenever SNP-outcome associations are estimated with varying degrees of precision and this is reflected in the analysis.

**Methods:** We propose instead to use a small modification of the scatter plot - the Galbraith radial plot - for the presentation of data and results from an MR study, which enjoys many advantages over the original method. On a practical level it removes the need to recode the genetic data and enables a more straightforward detection of outliers and influential data points. Its use extends beyond the purely aesthetic, however, to suggest a more general modelling framework to operate within when conducting an MR study, including a new form of MR-Egger regression.

**Results:** We illustrate the methods using data from a two-sample Mendelian randomization study to probe the causal effect of systolic blood pressure on coronary heart disease risk, allowing for the possible effects of pleiotropy. The radial plot is shown to aid the detection of a single outlying variant which is responsible for large differences between IVW and MR-Egger regression estimates. Several additional plots are also proposed for informative data visualisation.

**Conclusion:** The radial plot should be considered in place of the scatter plot for visualising, analysing and interpreting data from a two-sample summary data MR study. Software is provided to help facilitate its use.

## Background

Mendelian randomization (MR) [1] is a methodological framework for probing questions of causality in observational epidemiology using genetic data - typically in the form of single nucleotide polymorphisms (SNPs) - to infer whether a modifiable risk factor truly influences a health outcome. A particular MR study design gaining in popularity combines publically available data on SNP-exposure and SNP-outcome associations from separate but homogeneous studies for large numbers of uncorrelated SNPs. Each SNP is used to estimate the causal effect under the primary assumption that it is a valid instrumental variable (IV), by dividing its SNP-outcome association by its SNP-exposure association to yield the ratio estimate. Secondary modelling assumptions are also required in order for this estimate to be consistent. Ratio estimates are then combined into an overall estimate of causal effect using an inverse variance weighted (IVW) fixed effect meta-analysis. This is referred to as the IVW estimate and the general framework as two-sample summary data MR [2, 3]. For further details see Box 1.

#### Box 1: Standard two-sample summary data MR analysis

**The IV assumptions:** The canonical approach to MR assumes that group of SNPs are valid IVs for the purposes of inferring the causal effect of an exposure, *X*, on an outcome, *Y*. That is they are: associated with *X* (IV1); not associated with any confounders of *X* and *Y* (IV2); and can only be associated with *Y* through *X* (IV3). The IV assumptions are represented by the solid lines in the causal diagram below for a SNP *G*_*j*_, with unobserved confounding represented by *U*. Dotted lines represent dependencies between *G* and *U*, and *G* and *Y* that are prohibited by the IV assumptions. The causal effect of a unit increase in *X* on the outcome *Y*, denoted by *β*, is the quantity we are aiming to estimate.

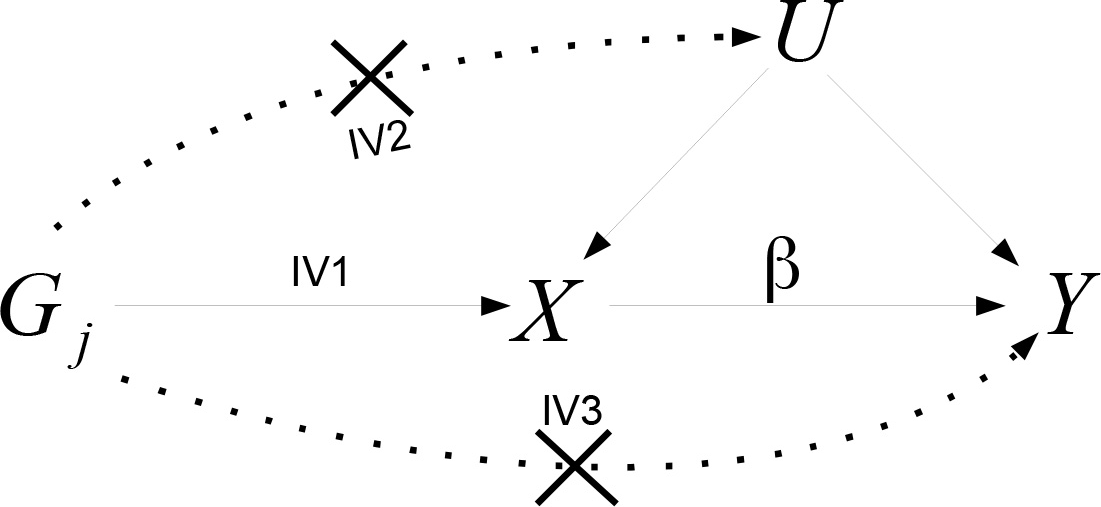

**The Ratio estimate:** Assume that exposure *X* causally affects outcome *Y* linearly across all values of *X*, so that a hypothetical intervention which induced a 1 unit increase in *X* would induce a *β* increase in *Y*. Suppose also that all *L* SNPs predict the exposure via an additive linear model with no interactions. If SNP *j* is a valid IV, and the two study samples are homogeneous, then the underlying SNP-outcome association from sample 1, Γ_*j*_, should be a scalar multiple of the underlying SNP-exposure association estimate from sample 2, γ_*j*_, the scalar multiple being the causal effect *β*. That is:

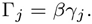

The ratio estimate for the causal effect of *X* on *Y* using SNP *j* (out of *L*), 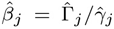, where 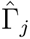 is the estimate for SNP *j*’s association with the outcome (with standard error *σ*_*Yj*_) and 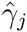 is the estimate for SNP *j*’s association with the exposure (with standard error *σ*_*Xj*_).

**The IVW estimate:** The overall inverse variance weighted (IVW) estimate for the causal effect obtained across *L* uncorrelated SNPs is then given by

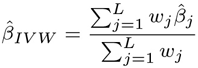

where *w*_*j*_ is the inverse variance of 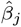. Two popular choices for the inverse variance weights are

1st order (fixed effect) weights: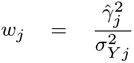
2nd order (fixed effect) weights: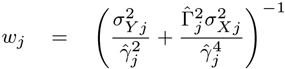

When SNP-exposure association estimates are sufficiently precise, so that 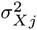 is negligible, or the causal effect *β* is small, then both weighting schemes are very similar. When this is not the case, both 1st and 2nd order weights can perform poorly. In this case, Bowden et al [4] propose the use of a ‘modified 2nd order’ weighting scheme instead.

Different formualae for the inverse variance weights can be employed, the most popular being simple ‘1st-order’ weights, which assume the uncertainty in the SNP-exposure association estimates is negligible. Although more sophisticated weighting approaches have recently been proposed [4], for simplicity we will use 1st order weights throughout this paper.

### The scatter plot

Figure 1 (left) shows a traditional scatter plot of summary data estimates for the associations of 26 genetic variants with systolic blood pressure (SBP, the exposure) and coronary heart disease (CHD, the outcome). SNP-exposure association estimates were obtained from the International Consortium for Blood Pressure consortium (ICBP) [5]. SNP-CHD association odds ratios were collected from Coronary ARtery Disease Genome-Wide Replication And Meta-Analysis (CARDIoGRAM) consortium [6], and then transformed to the log-scale for subsequent model fitting. These data have previously been analysed and interpreted by Lawlor et al [7] and Bowden et al [4]. They are included here for the purposes of illustration, rather than to draw any new epidemiological conclusions.

The ratio estimate for any individual variant is the slope joining its data point to the origin, as shown for a single variant in Figure 1 (left). The IVW estimate for these data, which represents the causal effect of a 1mmHg increase in SBP on the log-odds ratio of CHD, is 0.053. This is shown as the slope of a solid black line passing through the origin. The data point contributed by SNP rs17249754 is highlighted in red, as it will be subsequently discussed. It has become conventional to fix the sign of the SNP-exposure association estimates in these plots to be uniformly positive. This would naturally be achieved if each SNP had been coded to reflect the number of exposure increasing alleles. SNP-outcome association estimates must also be checked and altered to account for this change (see Box 2 for further details). This does not alter the result of the IVW analysis, but makes it easier to interpret the IVW estimate as a best fitting line through the data points.

**Figure 1:**
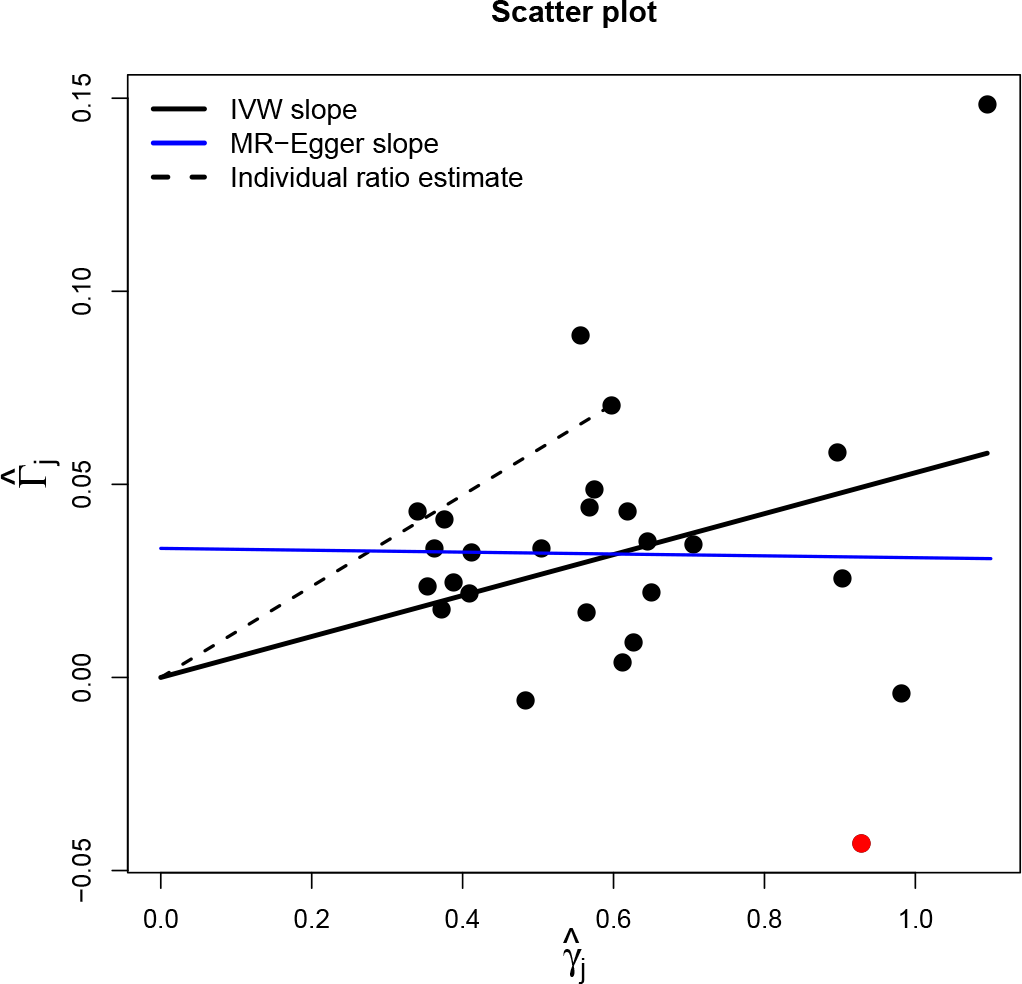
Traditional scatter plot of SNP-outcome associations 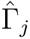 versus SNP-exposure associations 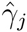, with IVW slope shown as a solid black line. SNP rs17249754 is highlighted in red.

##### Box 2: Detecting and accounting for heterogeneity in two-sample summary data MR

Heterogeneity amongst the ratio estimates can be calculated via Cochran’s *Q* statistic. When 1st order weights are used for the *w*_*j*_, *Q* can be expressed in two ways:

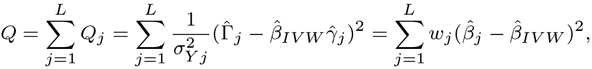

If heterogeneity is detected this suggests violation of the modelling or IV assumptions. Although horizontal pleiotropy is just one factor among many others that could be the underlying source of heterogeneity, we will assume it is the cause when explaining the implementation and assumptions of subsequent methods.

**Accounting for pleiotropy via a random effects meta-analysis:**. Let *α*_*j*_ equal the pleiotropic effect of SNP *j* on the outcome *Y* not through *X*, with sample mean and variance across all *L* SNPs of *μ*_*α*_ and 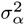 respectively. If *α*_*j*_ is independent in magnitude of the instrument strength across all SNPs (the InSIDE assumption) *μ*_*α*_ and *μ*_*α*_ = 0 (balanced pleiotropy), then an additive [24] or multiplicative [25] random effects meta-analysis can be used to reliably estimate the causal effect.

**Accounting for pleiotropy via MR-Egger regression:**. If *μ*_*α*_ is non-zero (directional pleiotropy) then the IVW estimate will generally yield a biased estimate for the causal effect. However if the InSIDE assumption holds then MR-Egger regression [11] can still deliver reliable estimates for the causal effect, along with an estimate for *μ*_*α*_. It is implemented by fitting the following linear regression of the SNP-outcome associations versus the SNP-exposure associations

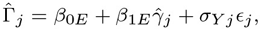

where ϵ_*j*_ ~ *N*(0,1)
after pre-processing the data according to the following rule:

For all *j* in (1,‥,L) such that 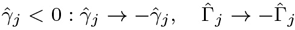.

The standard implementation of MR-Egger regression tacitly assumes 1st order weights. In this case, the InSIDE assumption is that the pleiotropic effects weighted by *σ*_*Yj*_ are independent of the SNP-exposure associations weighted by *σ*_*Yj*_.

**Assessing heterogeneity about the MR-Egger fit:** Heterogeneity about the MR-Egger fit can be assessed using Rücker’s *Q*′ statistic [12, 3]. When 1st order weights are used for the *w*_*j*_, *Q*′ can be expressed in two ways:

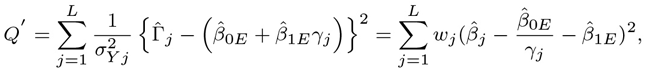

Specifically *Q*′ tests for the presence of heterogeneity due to pleiotropy around the MR-Egger fit after adjustment for its mean value, *μ*_*α*_ (estimated by 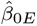). This is equivalent to testing whether 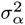 is greater than zero. When such heterogeneity is detected, standard errors for the MR-Egger intercept and slope parameter estimates, 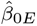 and 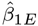 can be inflated by a factor of 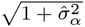. This is consistent with applying a multiplicative random effects model using 1st order weights.

### Detecting and adjusting for heterogeneity in two-sample MR

Within the meta-analytical framework underpinning the standard IVW estimate, heterogeneity observed amongst the ratio estimates can be assessed via Cochran’s *Q* statistic. If the necessary modelling assumptions hold for two-sample summary data MR and all SNPs are valid IVs, then Cochran’s *Q* should follow, asymptotically, a Chi-squared distribution, with degrees of freedom (df) equal to the number of SNPs minus 1. Excessive heterogeneity therefore points to a meaningful violation of some or all of these assumptions. Much attention has focused on detecting and adjusting for one specific source of violation referred to as *horizontal pleiotropy* [8, 9]. This occurs when SNPs exert a direct effect the outcome through pathways other than the exposure. For brevity we refer to this simply as ‘pleiotropy’ from now on. We will focus solely on this source heterogeneity for the rest of the paper, but return the topic of other sources of heterogeneity in the discussion.

Del Greco et. al. [10] first proposed the use of Cochran’s *Q* to detect pleiotropy in a Mendelian randomization context. However, the presence of heterogeneity due to pleiotropy does not automatically invalidate the IVW estimate. For example, if across all variants:

- (i) Its magnitude is independent of instrument strength (the so-called ‘InSIDE’ assumption [11]);
- (ii) It has a zero mean (i.e. it is ‘balanced’);

then a random effects meta-analysis can be used in lieu of the standard fixed effects IVW meta-analysis to reliably estimate the causal effect accounting for the additional uncertainty due to pleiotropy. If (i) holds but not (ii) then MR-Egger regression can instead be used to reliably estimate the mean directional pleiotropic effect and causal effect. [11, 3]. For the blood pressure data in Figure 1, and assuming pleiotropy is the source of heterogeneity, MR-Egger regression estimates the mean pleiotropic effect (i.e. the intercept) to be 0.033 and the causal effect adjusted for pleiotropy (i.e. the slope) to be virtually zero. Thus, MR-Egger infers that the effect detected by the IVW approach is spurious, and due to bias rather than any underlying causal mechanism.

An extended version of Cochran’s Q statistic (Rücker’s *Q*′ [12, 3]) can be used to assess heterogeneity about the MR-Egger fit. See Box 2 for further details. The size of *Q* and *Q*′ in relation to one another (specifically the difference *Q*-*Q*′) gives an indication as to the relative goodness of fit of the IVW and MR-Egger models. For this reason, Bowden et al [3] suggest reporting the statistic *Q*_*R*_ = *Q*′ /*Q* to aid the interpretation of study results from an MR-analysis. A *Q*_*R*_ close to 1 indicates the IVW and MR-Egger models fit the data equally well, whereas a *Q*_*R*_ much less than 1 indicates MR-Egger is best fitting. They also adapt the hierarchical model selection framework outlined by Rücker et al [12] for guiding which approach is appropriate for a given analysis. See Box 3 for further details. In essence this framework favours the use of the IVW model over MR-Egger regression *a priori* because it yields causal estimates with higher precision, but recommends MR-Egger regression only when it provides a demonstratively better fit to the data.

##### Box 3: The Rücker model selection framework

The Rücker model selection framework [12, 3] is encapsulated in the diagram below

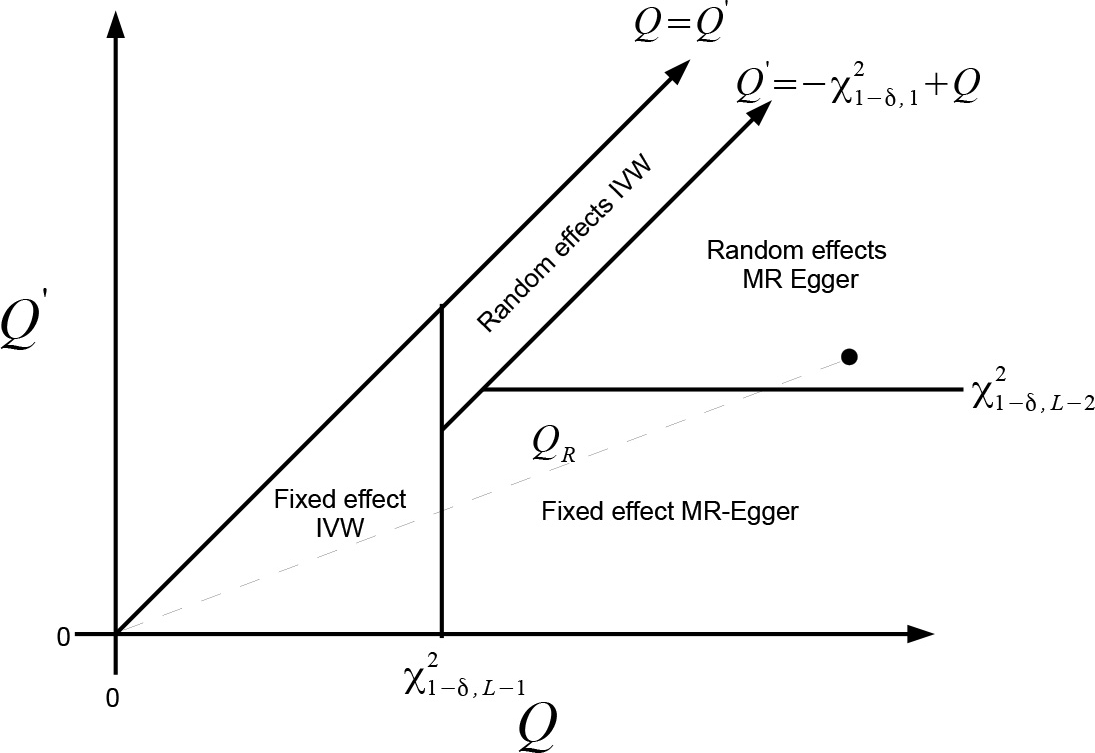

It shows the two-dimensional decision space defined by *Q*, *Q*′ and a significance threshold for detecting pleiotropy, δ (e.g. δ = 0.05). The rationale for this framework is briefly summarised.

1. Start by performing an IVW analysis under a fixed effect model and calculate *Q*.
2. If *Q* reveals sufficient heterogeneity at significance level δ with respect to a 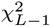 distribution then switch instead to a random effects IVW model.
3. Fit fixed effect MR-Egger regression and calculate *Q*′. If the difference *Q* − *Q*′ is significant at level δ with respect to a 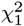 distribution, switch to this model.
4. If *Q*′ reveals sufficient heterogeneity at significance level δ with respect to a 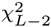 distribution then switch instead to a random effects MR-Egger model.

For a given data set, the slope joining the point (*Q*, *Q*′) to the origin gives the ratio statistic *Q*_*R*_,and the point (*Q*, *Q*′) immediately defines the selected model under the above framework. This is illustrated by the black dot in the diagram above. In this hypothetical case the Rücker framework suggests the random effects MR-Egger model is most appropriate [3].

Aligning the SNP-exposure association estimates is irrelevant to IVW analysis since the IVW estimate remains constant whichever coding is used. However, it is actually a necessary step for the standard implementation of MR-Egger regression. This can be understood by viewing MR-Egger as a method for detecting and adjusting for any systematic trend in the causal estimates according to the ‘weight’ each one receives in the IVW analysis, with weight being a strictly positive quantity.

### Limitations of the scatter plot for MR analysis

Although it has become the standard tool for visualizing summary data in an MR analysis, the scatter plot has a major limitation, which lies at the heart of this paper:

The scatter plot does not give the most transparent representation as to the weight each genetic variant receives in the MR analysis, whenever the weights are not solely determined by the SNP-exposure associations.

**Figure 2:**
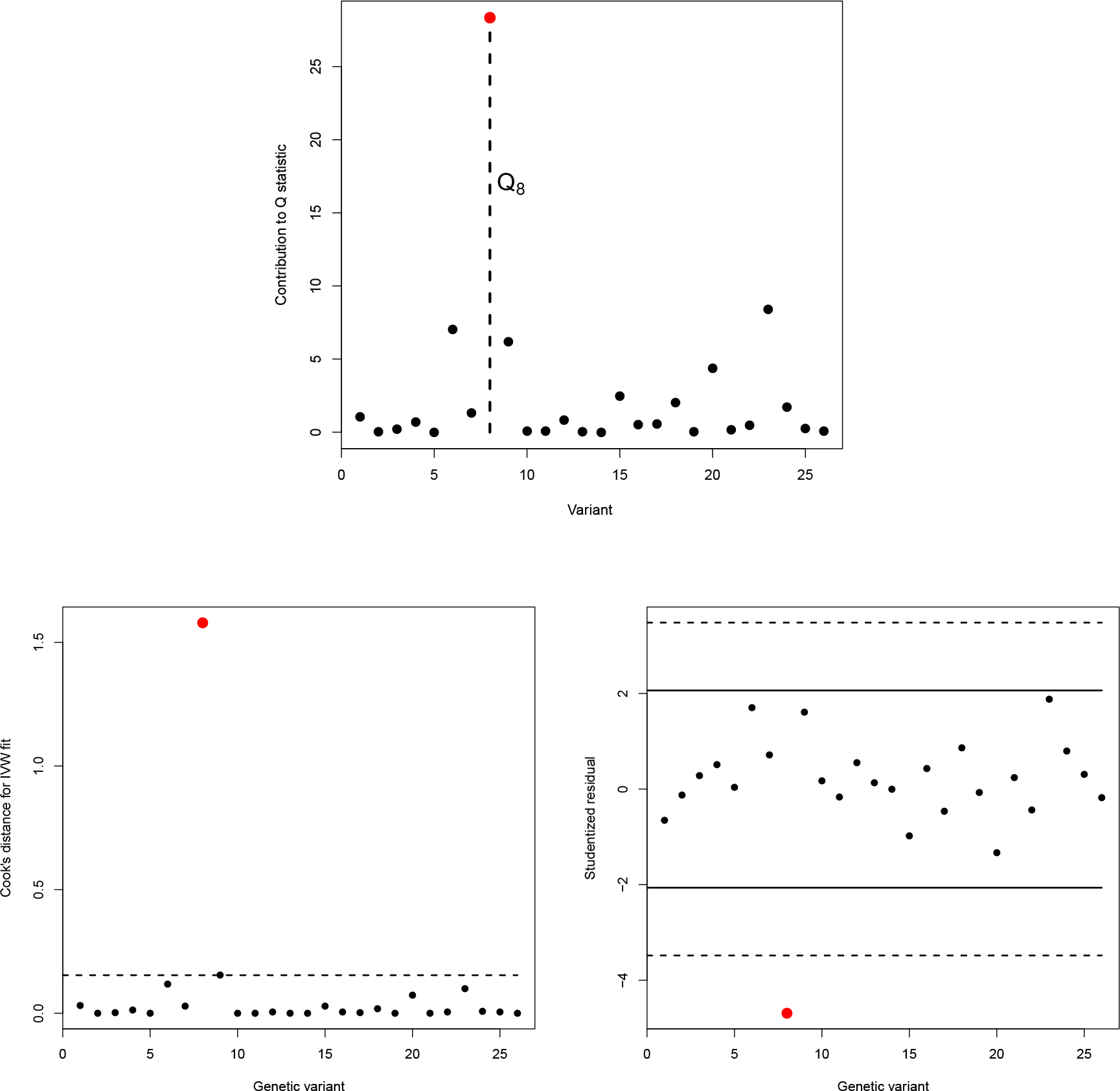
Top: Individual variant contributions to Cochran’s heterogeneity statistic. The contribution of SNP rs1724 9754 (labelled Q_8_) is highlighted in red. Bottom left: Cook’s distance for each genetic variant in the SBP-CHD data, with standard influence threshold (4/#SNPs) indicated by a dashed line. Bottom right: Studentized residuals for each variant in the SBP-CHD data with standard 5% signifance thresholds (solid black lines) and bonferroni corrected significance thresholds (5%/#SNPs, dashed lines). SNP rs1724 9754 is again shown in red.

This is the case for the IVW estimate calculated using standard 1st order weighting, and shown as a fitted slope in Figure 1, since they depend additionally on the SNP-outcome association standard error. This lack of transparency hampers the visual detection of outliers and influential data points in the analysis, for example SNP rs17249754 highlighted in red, which is illustrated further in Figure 2. In Figure 2 (top) we plot the value of each individual variant’s contribution to Cochran’s *Q* statistic, which is approximately Chi-squared distributed with 1 df under the previously stated assumptions. For these data *Q* = 67.09 (df = 25), indicating substantial heterogeneity, but the individual contribution of SNP rs17249754 (the eighth variant in our data frame highlighted in red) is 28.34. It is therefore responsible for the vast majority of excess heterogeneity amongst the 26 ratio estimates.

Figure 2 (Bottom-left and-right) shows the Cook’s distance and Studentized residual measures for each variant, that were first used by Corbin et. al. [13] to look for influential SNPs in an MR context. Both measures also confirm rs17249754 as *the* major outlier for these data. However, this fact would not be immediately obvious from a visual inspection of the scatter plot alone.

## Methods

### The Radial MR plot

The Galbraith radial plot [14, 15] was proposed as a graphical tool to visualise estimates of the same quantity with varying precisions. Specifically it plots the *Z*-statistics for each estimate (i.e. the point estimate divided by its standard error) on the vertical axis versus the inverse standard error on the horizontal axis. It has been used extensively in meta-analysis to detect heterogeneity and small study bias [12, 16, 17]. We believe that, when translated to the MR-setting, it offers a simple solution to the inherent deficiencies of the standard scatter plot. The horizontal axis of the radial plot is the square root of the actual inverse variance weight each SNP receives in the IVW analysis. Its vertical axis scale represents the ratio estimate for each SNP multiplied by the same square root weight. Since the square root weight on the horizontal axis is naturally positive, and the vertical axis is a function of this same weight and the ratio estimate (which is coding invariant), the radial plot removes the need to manually re-orient the summary data estimates. Figure 3 (left) shows the blood pressure data, this time represented on the radial MR plot. The IVW estimate is again overlaid on top.

The radial plot still enables the slope joining each data point to the origin to be interpreted as a ratio estimate. A second vertical axis is usually drawn on the right hand side of the radial plot as an arc to accentuate this point. We leave this out in this instance in order to focus attention on the new scale of the horizontal and vertical axes only. An additional helpful property of the radial plot is that the absolute vertical distance from each data point to the fitted IVW slope is equal to the square-root of its contribution to Cochran’s *Q* statistic. From the radial plot we can instantly see that SNP rs17249754 is the most influential variant in the IVW analysis for two reasons:

a. It gets the most weight because of its position on the horizontal axis;
b. It has the largest contribution to Cochran’s *Q* statistic because it is furthest away from the IVW slope.

**Figure 3:**
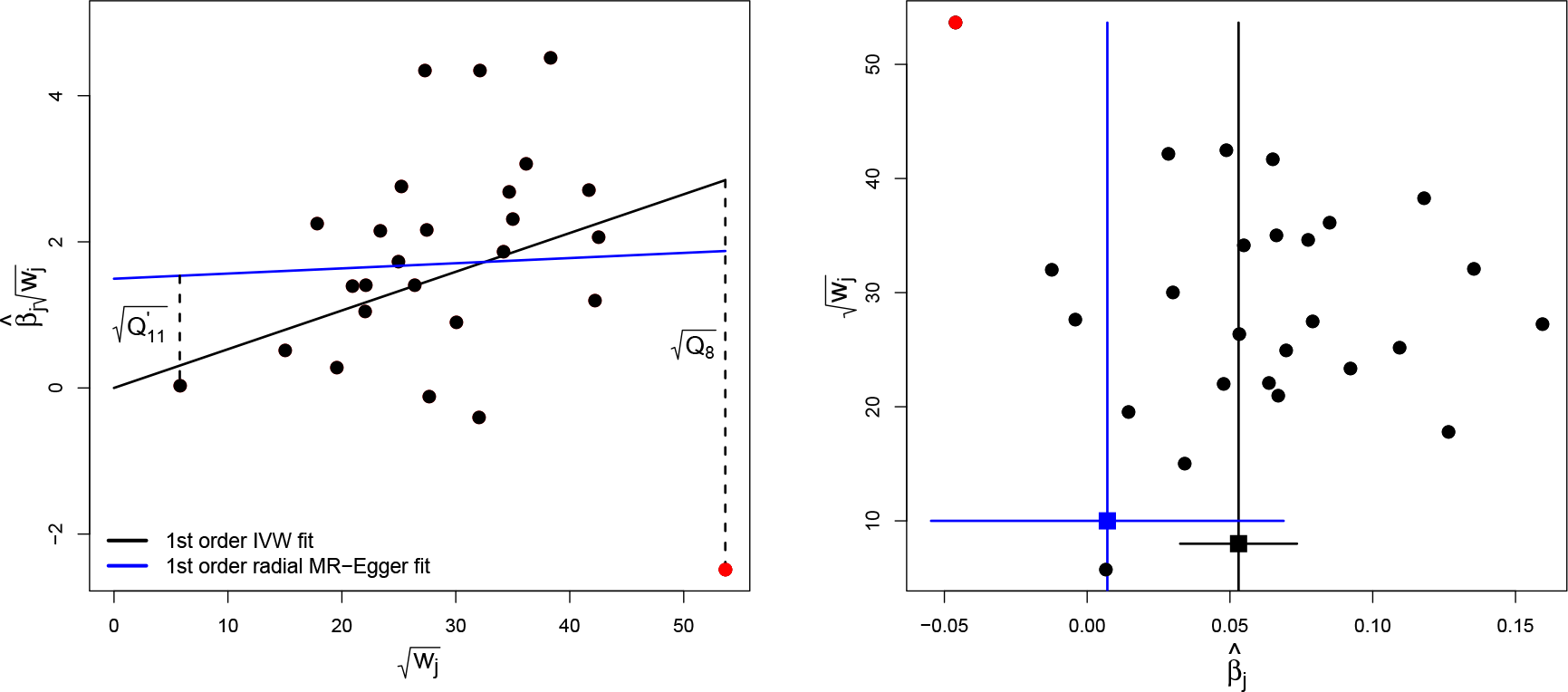
Left: Radial MR plot of the blood pressure data. IVW and radial MR-Egger regression slopes calculated using 1st order weights are overlaid. The square root contribution of SNP rs17249754 to Cochran’s Q statistic 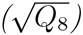 is denoted by the vertical dashed line from the IVW slope. The square root contribution of a separate SNP to Rücker’s Q′ statistic 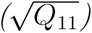 is denoted by the vertical dashed line from the radial MR-Egger slope. Right: Generalized funnel plot of same data with 1st order IVW and radial MR-Egger regression slopes (and 95% confidence intervals) shown. SNP rs1724 9754 is highlighted red.

### MR analysis via radial regression

Although the standard meta-analysis formula can be used to derive the IVW estimate (Box 1), in practice it is often convenient to obtain the estimate by fitting a linear regression model. This is a simple command in any software package, and allows the user to benefit from the host of summary and diagnostic tools that compliment it. For example, regressing the SNP-outcome associations on the SNP-exposure associations with the intercept constrained to zero, and weighting the regression by the SNP-outcome association standard error will yield the IVW estimate using 1st order weights. More generally, we can interpret the IVW estimate calculated using *any* set of user-defined weights as a best fitting line through the data points on the radial plot under the constraint that the line goes through the origin. See Box 4 for further details.

##### Box 4: Two-sample summary data MR via radial plot regression

**Radial IVW regression:** The IVW estimate obtained using any set of weights *w*_*j*_ can be interpreted as the *β* coefficient estimated from the following IVW Radial regression model:

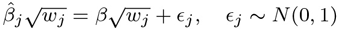

Cochran’s *Q* statistic must then be calculated as

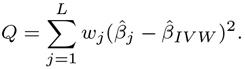

**Radial MR-Egger regression:** As a natural compliment to the radial IVW model above, the following radial MR-Egger regression model below can instead be used to estimate the causal effect:

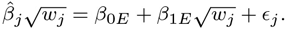

That is, radial MR-Egger is a regression directly on the radial plot scale with the intercept parameter left unconstrained. Under a radial model the InSIDE assumption is that the pleiotropic effects are independent of the radial weights.

Rücker’s *Q*′ statistic for the radial MR-Egger model is defined as:

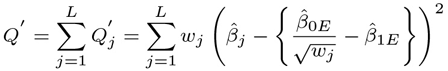

Fixed effect and random effects versions of radial IVW and radial MR-Egger regression can be implemented by altering the definition of *w*_*j*_.

**How does this differ from traditional MR-Egger?** The originally proposed MR-Egger regression model, which implicitly used 1st order weights, is equivalent to the following radial MR-Egger regression model:

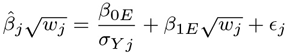

where *w*_*j*_ represent 1st order weights. That is, *β*_*0E*_ in the original model is not a true intercept (i.e. a constant), it is the coefficient of the explanatory variable 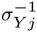, as explained in [3]. In practice traditional and radial MR-Egger will yield qualitatively similar inferences, although the magnitude of their respective intercept parameters will be different.

**Generalised radial plots** A generalised funnel plot that naturally complements the radial plot can be produced by plotting 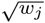 on the vertical axis against 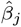 on the horizontal axis. This plot, however, is most nformative for the IVW analysis because the IVW slope lies at the (inverse variance weighted) centre of the data points. An equivalent radial MR-Egger funnel plot with the same property can be produced by plotting 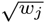 on the vertical axis against

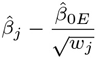

on the horizontal axis.

Just as for the IVW estimate, MR-Egger regression can also be implemented as a linear regression directly on the radial plot, but with the intercept left unconstrained. We call this *radial MR-Egger regression*. Radial MR-Egger regression is different from traditional MR-Egger regression, even when 1st order weights are used, because the intercept parameter is estimated on a different scale. Estimates obtained from a radial MR-Egger regression will be consistent for the causal effect as long as the InSIDE assumption is satisfied on this new scale (see Box 4).

Figure 3 (left) shows the radial MR-Egger regression slope, estimated assuming 1st order weights. Just as for the IVW method, the absolute distance from any data point to the radial MR-Egger slope is equal to the square root of its contribution to the overall heterogeneity after adjustment for pleiotropy - which is measured for MR-Egger by Rücker’s *Q*′ statistic. This is illustrated in Figure 3 for a single SNP. Note that the definition of Riicker’s *Q*′ also changes under this analysis (Box 4). The radial plot can therefore be used to simultaneously assess whether individual variants are outliers with respect to either the IVW or radial MR-Egger regression models.

### Generalized funnel plots

Figure 3 (right) shows the blood pressure data represented on the funnel plot. It plots the ratio estimate for each variant on the horizontal axis against its square root precision (or weight) on the vertical axis. In this instance, 1st order weights were used to scale the vertical axis and to calculate the IVW and radial MR-Egger regression slope estimates which are overlaid on top. Under 1st order weighting, Figure 3 (right) is equivalent to the funnel plot first used by Bowden et al [11] to visualise MR analyses, and to look for asymmetry as a sign of pleiotropy. However, we label the vertical axis generically to stress that a generalized funnel plot can be produced, and will naturally compliment its corresponding radial plot, when any given set of weights are used.

Although it is possible to interpret the radial plot simultaneously for IVW and radial MR-Egger regression, the funnel plot in Figure 3 (right) is predominately informative about the IVW analysis. Specifically, the IVW estimate intuitively lies in the ‘centre of mass’ of the data when the mass of each ratio estimate is equated with its weight. This is explained in detail by Bowden and Jackson [18]. In order to produce a funnel plot with this same property for radial MR-Egger, we must apply a transform to the ratio estimate of each data point in the funnel plot, by subtracting the radial MR-Egger intercept estimate divided by the ratio estimates’ square root weight [18] (see Box 4). This is shown by the horizontal dashed lines in Figure 4. Because it is inversely proportional to the square root weight, the correction will be larger for imprecise ratio estimates and smaller for precise estimates. The correction factor for the least precise (11th) ratio estimate, 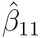 is explicitly labeled. We can relate and cross-reference this to the corresponding radial plot in Figure 3 (left), where the 11th ratio estimate is also labelled. It is not an outlier in the IVW analysis because of its proximity to the IVW slope, but its distance from the radial MR-Egger slope is far greater.

**Figure 4:**
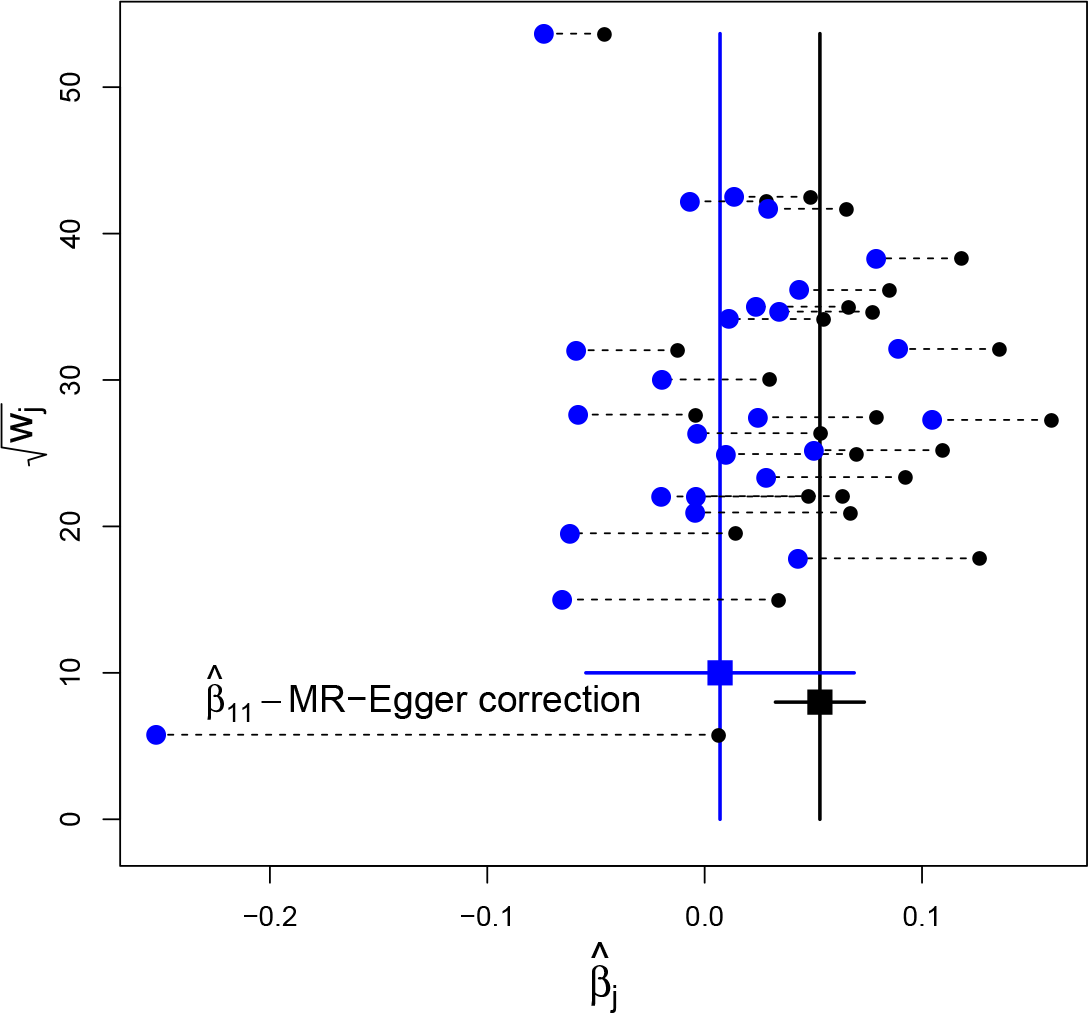
Left: Radial MR-Egger funnel plot. Horizontal dashed lines link the position of data in the standard funnel plot (black) to their implied position under a radial MR-Egger analysis (blue).

## Results

Table 1 shows results of our re-analysis of the blood pressure data using IVW and MR-Egger regression, first with all 26 SNPs and then with SNP rs17249754 removed. For comparison we show results for both the standard and radial implementation of MR-Egger regression. All analyses were carried out using 1st order weights, and assuming a multiplicative random effects model if any residual heterogeneity was detected.

The IVW estimate for the causal effect of a 1mmHg increase in SBP on the log-odds ratio of CHD is 0.053. Large heterogeneity is present amongst the 26 ratio estimates, as identified by Cochran’s *Q*, which is sufficiently extreme (p=1×10^−5^) to opt for a random effect IVW model instead. Standard and radial MR-Egger regression yield qualitatively similar results and suggest a causal effect close to zero. Both models represent a better fit to the data at well below the conventional 5% threshold since in each case *Q* - *Q*′ is much larger than 3.84 (the 95 percentile of a Chi-squared distribution on 1 df). Since a large amount of residual heterogeneity was still present around both the standard and radial MR-Egger fits (as detected by *Q*′), their standard errors were also inflated to allow for overdispersion.

When the three analysis methods are repeated this time with variant rs17249754 removed, IVW and MR-Egger causal estimates are virtually identical, especially with those of radial MR-Egger. Cochran’s *Q* and Rücker’s *Q*′ statistic only reveal a small amount of residual heterogeneity and examination of *Q* - *Q*′ reveals neither standard nor radial MR-Egger represent a better fit to the data than the IVW model. Therefore, the data does not support a move away from the standard IVW analysis without SNP rs17249754.

**Table 1:**
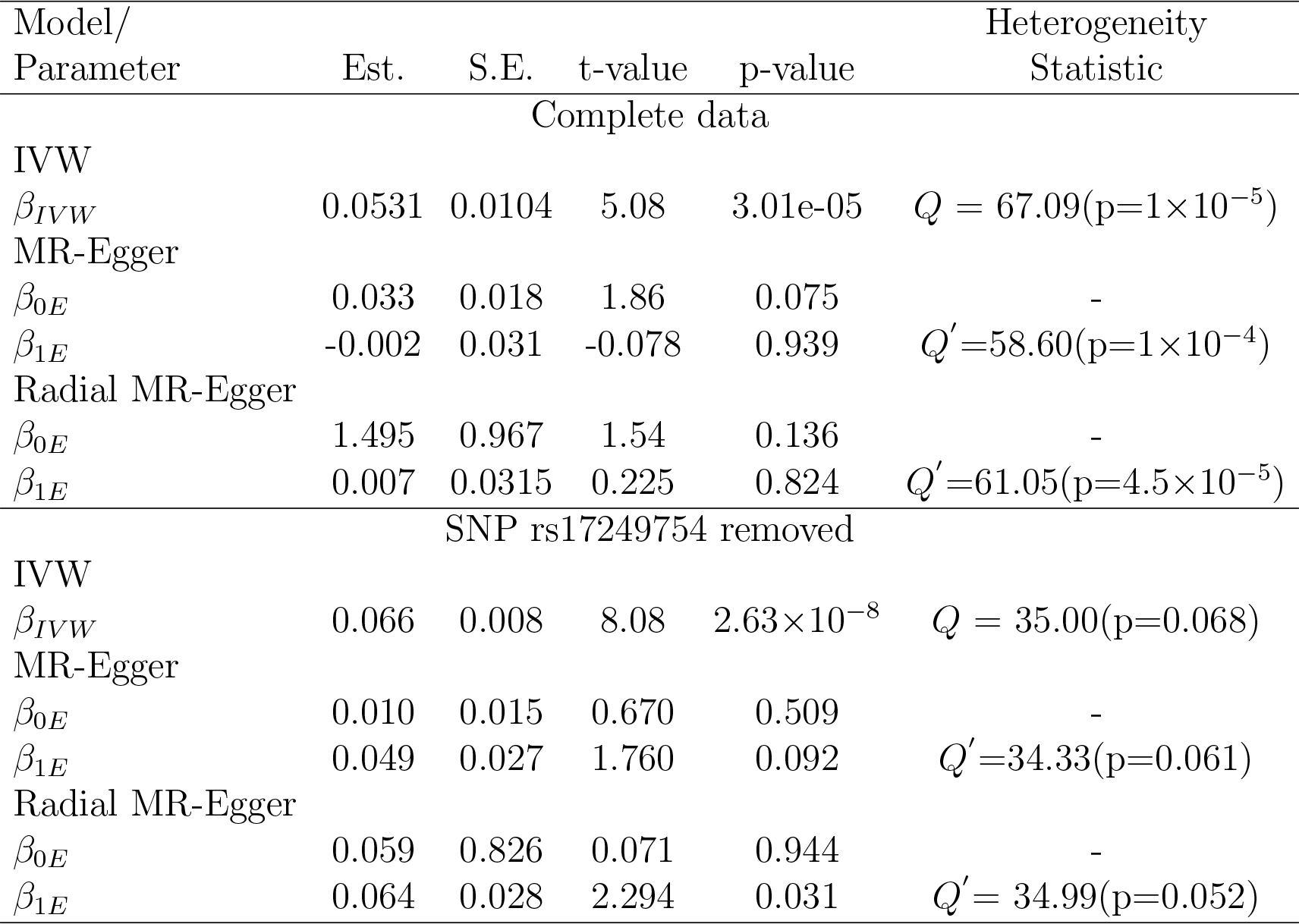
IVW and MR-Egger regression analyses of the SBP data with all SNPs and with SNP rs1724 9754 removed. Multiplicative random effects models were fitted in all cases whenever over-dispersion was detected.

### A leave-one-out sensitivity analysis

Rather than using the Rücker framework for formal model selection purposes (Box 3), we instead demonstrate its utility in providing a useful, but informal, backdrop to assess the influence of each individual variant on the analysis under the IVW and MR-Egger frameworks. Figure 5 shows the values of Cochran’s *Q* (calculated with respect to the IVW fit) against Rücker’s *Q*′ (calculated with respect to the radial MR-Egger fit) for 26 analyses where each SNP is left out in turn. These points are overlaid on top of the Rücker decision space assuming a threshold of δ = 0.05 for declaring heterogeneity using *Q* and *Q*′. In the main analysis reported in Table 1, random effects models were fitted if *any* heterogeneity at all was detected, which is equivalent to setting δ = 0.5. The nested nature of the radial IVW and MR-Egger models guarantees that all points in Figure 5 lie below the diagonal line *Q* = *Q*′.

When all the data are analysed together (orange point in Figure 5), sufficient heterogeneity and bias are detected to mean that a random effects radial MR-Egger regression model is best supported by the data. It infers the presence of large directional pleiotropy and no causal effect between SBP and CHD risk. This is not materially changed when every variant *except* SNP rs17249754 is left out of the analysis in turn (black dots in Figure 5). However, when SNP rs17249754 is removed from the data (red dot in Figure 5), there is no evidence of heterogeneity or bias due to directional pleiotropy and the data provides no reason to move away from a standard IVW analysis.

**Figure 5:**
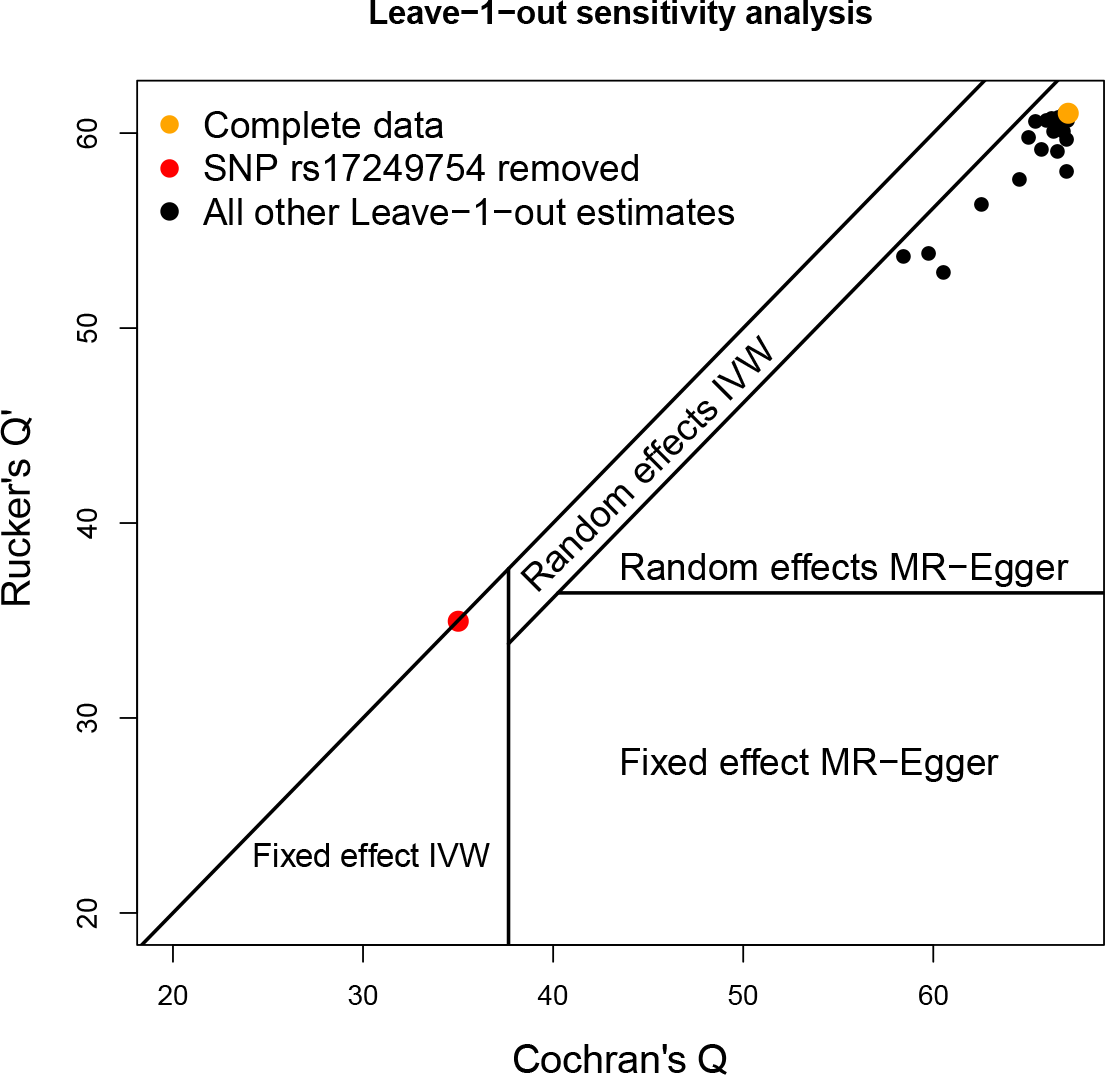
Leave-one-out sensitivity analysis of the data, showing the values of Q and Q′ when each variant is left out of the analysis in turn. Points are overlaid on the Rücker decision space that governs which of four model choices should be favoured. It assumes a significance threshold of δ = 0.05 to affect the model selection.

### The radial plot function

We have written an R function RadialMR to produce radial plots and to perform radial regression. Two of the many possible plot options are illustrated for the blood pressure data in Figure 6. Figure 6 (top) shows the radial plot of the IVW analysis alone, which includes a radial curve to highlight the ratio estimate for each genetic variant, as well as the overall IVW estimate. Data points with large contributions to Cochran’s Q statistic are shown in orange. The significance level for identifying these outliers can be set by the user, here we chose the value 0.01. Figure 6 (bottom) shows the radial plot on a tighter scale, with both IVW and radial MR-Egger regression implemented. Outliers for either method (and both methods) are shown. A table of the exact *Q* and *Q*′ contributions for each variant is given as an output for the researcher to conduct a more detailed analysis.

Radial plots are produced by many existing R packages such as `metafor`, `numOSL`, and `Luminescence`. Care will need to be taken, however, to input data from an MR-analysis appropriately into these generic platforms. For this reason we will also continue to develop our own `RadialMR` package to produce radial plots and conduct radial plot regression for the MR-setting. It is currently available to download at https://github.com/WSpiller/RadialMR/ and will continue to be refined and extended to incorporate further features and analyses.

## 1 Conclusion

It has long been appreciated in the general meta-analysis context that the radial plot has many desirable characteristics over the traditional scatter plot, especially in the detection of outlying studies and small study bias. Given its intimate connection with meta-analysis, we propose that the radial plot should also be given a more central role in two-sample summary data MR studies.

The radial plot, and its corresponding funnel plots, improve the visual interpretation of data used within an MR analysis because it provides the most transparent representation from an information content perspective. Its implications stem beyond the purely aesthetic for MR-Egger regression, however. Radial MR-Egger is an attractive modification and generalization of the original approach that naturally flows from the use of this plot. On top of removing the need to recode the genetic data and facilitating a more straightforward detection of outliers, the radial formulation also makes it much more transparent that it is attempting to detect any systematic trend in ratio estimates according to the weight they receive in the analysis. Another advantage is that it only requires the ratio estimates and their standard errors. This makes it applicable even when data on individual SNP-exposure and SNP-outcome associations (and their standard errors) are not available.

**Figure 6:**
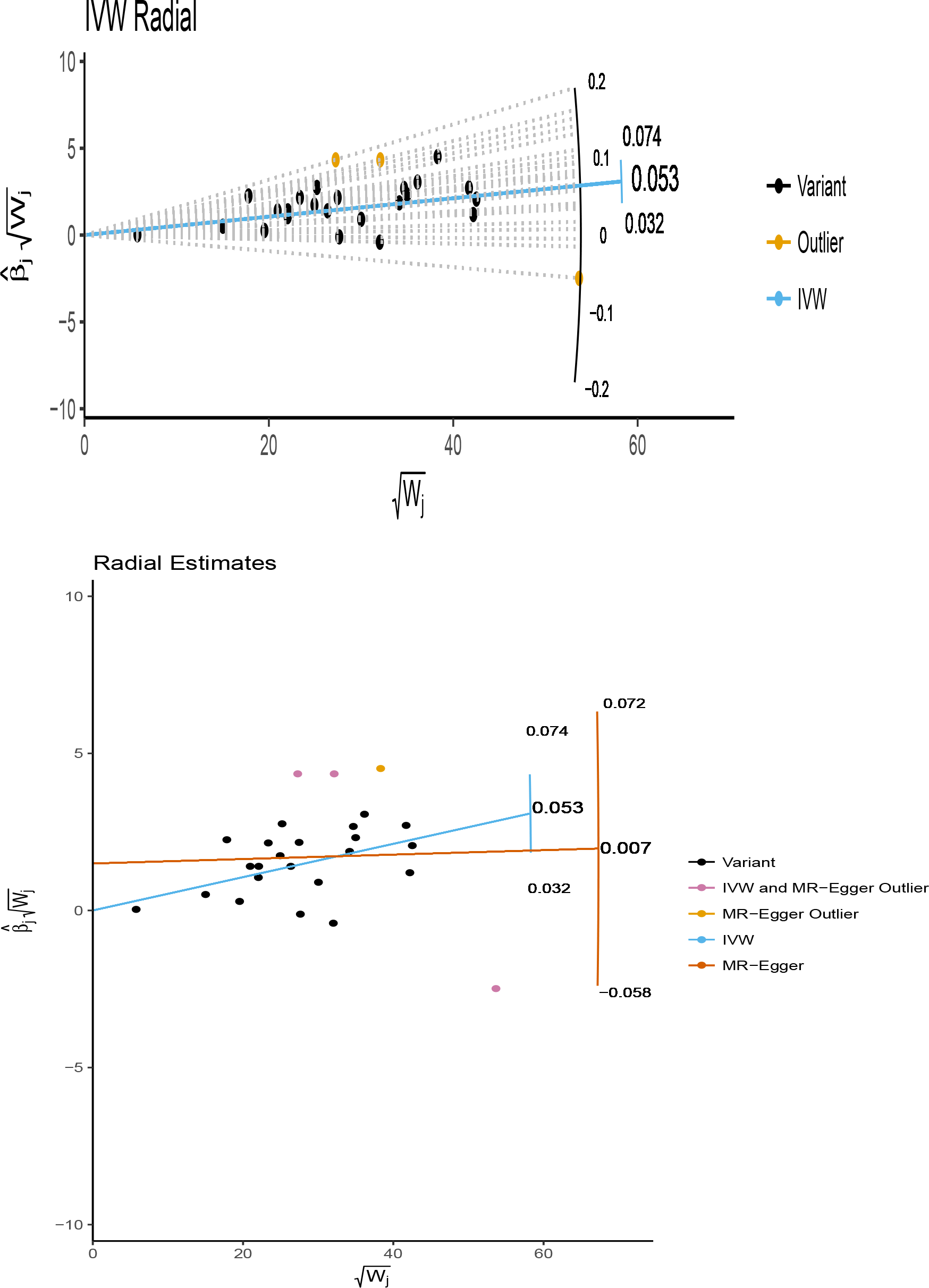
Radial plots of the blood pressure data produced using the RadialMR() function. Top: only the IVW estimate shown, radial lines joining each data point back to the origin. Bottom: Radial MR-Egger andJVW model fits shown.

When 1st order weights are used, radial MR-Egger and traditional MR-Egger will yield similar causal estimates, but the magnitude of the intercept will be different. An undoubted strength of the radial approach lies in the fact that it can be seamlessly applied when any set of weights are used. For example, Bowden et al [4] have shown that 1st order weights can inflate the type I error rate of Cochran’s *Q* and Rücker’s *Q*′ statistics for detecting heterogeneity, when none is present, whenever the SNPs utilized are collectively weak or there is a large causal effect. They propose modified weights that depend on the causal estimate to improve the performance of both statistics and those of the IVW and MR-Egger estimates. These weights (and indeed any weights) can be naturally incorporated into a radial plot, the IVW approach and radial MR-Egger. Further investigation into the properties of radial MR-Egger in a variety of circumstances are required, but the features that distinguish it from the standard approach appear attractive, and it has the potential to become the standard implementation.

When conducting a two-sample summary data MR analysis with a binary outcome, natural correlations will exist between causal effect estimates (e.g. log-odds ratios) and their precisions, which could easily contribute to heterogeneity and hence be misconstrued as pleiotropy. In related work on the meta regression of separate trial results measuring a binary outcome, Harbord et al [19] show that regressing the ratio of the score and square root information statistics against the square root information (in a close analogy to the radial plot) is better at mitigating this effect than simply working directly with the log-odds ratio and its standard error. As further work we plan to extend the approach of Harbord to the MR context for radial MR-Egger regression with binary outcomes. Similar approaches based on score and information statistics may also prove useful for MR analyses of time-to-event outcomes.

We have proposed a leave-one-out analysis using the Rücker model selection framework as a backdrop when conducting an MR study, to understand how model choice is affected by the exclusion of individual variants. However, we stress some caution in following this approach to the extreme, for example in adopting a strategy of removing multiple outliers until little or no-heterogeneity remains, unless its statistical properties are well understood. Procedures such as this have been proposed when meta-analysing separate study results [20], but have been criticised for being too data driven, likely to throw out larger studies than smaller studies, and offering little explanation as to the underlying cause of heterogeneity [21].

The Rücker model selection framework we present explores how the choice of IVW or MR-Egger model is affected by the summary data from each SNP, but it can not tell the user about the probability each model is true. Thompson et al [22] have proposed a formal Bayesian model averaging framework that achieves this aim, and produces posterior causal effect estimates accounting for model uncertainty. Hemani et al [23] have also recently proposed a machine learning framework for choosing between a much larger group of modelling choices. Both ideas nicely compliment and extend the basic the approach outlined here.

#### Key messages

- Summary data furnishing a two-sample Mendelian randomization study are often visualized with the aid of a scatter plot. The scatter plot is also used to interpret the validity of the standard inverse variance weighted (IVW) estimate, and pleiotropy robust methods such as MR-Egger regression.
- A close relation of the scatter plot - the radial plot - can instead be used for this purpose.
- The radial plot removes the need to pre-process the summary data (a prerequisite for MR-Egger), improves the detection of outliers and influential data points in either an IVW or MR-Egger analysis, and can incorporate any set of weights desired by the user.
- A more general form of MR-Egger regression is proposed that flows from, and naturally compliments, the radial plot.
- Generalized funnel, and leave-one-out analysis plots can also be used to aid the visualisation and interpretation of MR studies.

